# Software update: Moving the R package *sommer* to multivariate mixed models for genome-assisted prediction

**DOI:** 10.1101/354639

**Authors:** Giovanny Covarrubias-Pazaran

## Abstract

In the last decade the use of mixed models has become a pivotal part in the implementation of genome-assisted prediction in plant and animal breeding programs. Exploiting the use genetic correlation among traits through multivariate predictions has been proposed in recent years as a way to boost prediction accuracy and understand pleiotropy and other genetic and ecological phenomena better. Multiple mixed model solvers able to use relationship matrices or deal with marker-based incidence matrices have been released in the last years but multivariate versions are scarse. Such solvers have become quite popular in plant and animal breeding thanks to user-friendly platforms such as R. Among such software one of the most recent and popular is the *sommer* package. In this short communication we discuss the update of the package that is able to run multivariate mixed models with multiple random effects and different covariance structures at the level of random effects and trait-to-trait covariance along with other functionalities for genetic analysis and field trial analysis to enhance the genome-assisted prediction capabilities of researchers.

## Introduction

Currently, linear mixed models play an important role in science to better understand different biological and non-biological phenomena (Bolker 2009; Gianola and Rosa 2015). In brief, linear mixed models are extensions of linear models (Hastie et al., 2009). Among many statistical tools used to test hypothesis and estimate parameters, linear mixed models are particularly robust and flexible, which has given a pivoting role in
Biological sciences such as plant and animal breeding and ecology (Bolker et al. 2009; Hadfield 2010; Gianola and Rosa 2015). For example, in the Genetics field, most of the published genome wide association and quantitative genetics studies are based on mixed models and REML estimation (Bush et al. 2012; Hirschhorn et al. 2005; Wang et al.2005; Kang et al. 2008, 2010).

In general terms, current mixed model solvers use Frequentist or Bayesian approaches to estimate parameters and solve the linear equations, and both approach rely on statistical assumptions (i.e. distributions) on the response or covariates to be modeled (Gianola and Rosa 2015; Hastie et al., 2009). Among Frequentist software and different optimization techniques, mixed models are usually solved by either of the two most popular REML/ML methods; mixed model equation-based (MME) algorithms, based on Henderson and Searle ideas (Gilmour et al 1995; Henderson 1975; Searle 1993) and direct-inversion-based (DI) algorithms which is the natural solution for the linear equations in the mixed model context (Lee et al 2016; Maier 2015). Both methods require iterative procedures to estimate the variance-covariance parameters and coefficients given that closed solution does not exist when the variance-covariance parameters are not known and more than one variance component other than error needs to be estimated. The former method (MME-based) is particularly advantageous when the number of parameters to estimate is less than the number of observations (n > p scenario), whereas the latter method performs better when the number of parameters is greater than the number of observations and dense covariance structures are used (p > n). On the other hand, Bayesian approaches implement Markov chain Monte Carlo routines (carrying a large computational overhead), which in addition, require the specification of prior distributions, in particular for variance components (Fong et al. 2010). Numerous literature about these topics is available and a good source for history and sources to look at can be found in Gianola and Rosa (2015) and other textbooks.

Univariate mixed models have become very popular in multiple research fields thanks to the advent of open-source and easy to use software (Bates et al. 2015). On the other hand multivariate mixed models have been slowly adapted due to the limited open-source resources. In the biology fields, multivariate mixed models were popularized in animal breeding to model the genetic correlation among traits, longitudinal data and to model genotype by environment interactions (trajectory across multiple years or environments) (Mrode, 2014; Lee and van der Werf, 2016). Extensive animal breeding literature using multi trait models has been generated using multi trait pedigree-based BLUP (MPBLUP) (Schaeffer, 1984; Thompson and Meyer, 1986; Mrode, 2014). Currently, with the massive availability of genetic markers, multi trait models using genomic information (MGBLUP) are now feasible to model plant populations for crops where pedigree information is not robust and is becoming more and more popular given the significant increase in predictive ability (PA) that the use of this method brings in genomic selection (GS) programs (Calus and Veerrkamp, 2011; Jia and Janninck, 2012; Mrode, 2014; Marchal et al. 2016). Besides the use in GS programs multivariate models are very valuable in Ecology, Evolution and other fields (Hadfield et al. 2007). The availability of reliable open-source software can boost the implementation of such methods for the benefit of multiple research fields.

Within the open-source platforms, the R language is becoming increasingly popular in the statistical and biological communities. The existence of numerous libraries provides researchers the ability to use state of art techniques in their own analysis (R Core Team, 2017). Unfortunately, the availability of multivariate mixed model solvers in R has been limited. The most popular R packages available for this purpose are *ASReml-R* (licensed package) and*MCMC glmm* (Gilmour et al. 1995, Hadfield et al. 2010). Each of these packages has its own strengths and limitations. In this paper, we present the updated version of the *sommer* package, once released as a univariate mixed model solver for genome assisted prediction (Covarrubias-Pazaran, 2016). The new version of the *sommer* package has evolved to a high degree of sophistication to fit multivariate models with multiple random effects with different covariance structures at the level f the random effects and at the level of the trait-to-trait structures based on the Newton Raphson optimization method using direct inversion (DI) which is optimal for genome assisted prediction where the number of effects to estimate is bigger than the number of observation (p > n; Lee et al. 2006). In the following section we present a comparison with the most popular software and discuss the strengths and limitations of *sommer* against the available software and we propose the use of *sommer* as a complement to available software for the p > n scenario.

## Materials and Methods

### Multivariate algorithm

In order to face with the dimensionality issue with the advent of massive genomic information, defined as the p>n problem (more random and fixed effects parameters to estimate than observations; for example genetic markers), we decided to focus the software in the direct-inversion algorithms, which has been found to be faster than MME-based algorithms when p > n or when dense covariance structures are used (Lee and Vander Werf 2006). We coded the multivariate Newton-Raphson algorithm for multiple random effects and covariance structures in the updated version of the R package *sommer* available at the CRAN (https://CRAN.R-project.org/package=sommer). Following Maier et al. (2015), the multivariate mixed model implemented has the form:

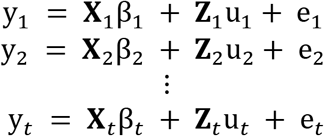

where y_i_ is a vector of trait phenotypes, β_i_ is a vector of fixed effects, u_i_ is a vector of random effects for individuals and e_i_ are residuals for trait ‘i’ (i = 1, …, t). The random effects (u_1_ … u_i_ and e_i_) are assumed to be normally distributed with mean zero. **X** and **Z** are incidence matrices for fixed and random effects respectively. The distribution of the multivariate response and the phenotypic variance covariance (**V**) are:

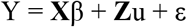
where 
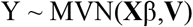

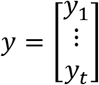

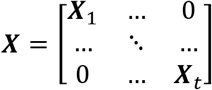

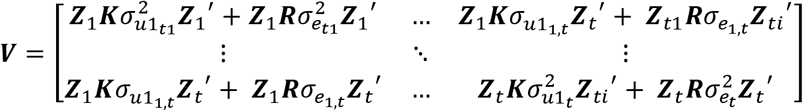

where **K** is the relationship or covariance matrix for the k^th^ random effect (u=1,…,k), and **R**=**I** is an identity matrix for the residual term. The terms, 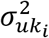 and 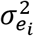 denote the genetic (or any of the k^th^ random terms) and residual variance of trait ‘i’, respectively and *σ*_*uk*_*ij*__and *σ*_*e*_*ij*__ the genetic (or any of the k^th^ random terms) and residual covariance between traits ‘i’ and ‘j’ (i=1,…,t, and j=1,…,t). The algorithm implemented optimizes the log likelihood:

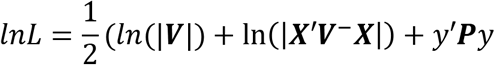

where ln is the natural log, and | | the determinant of the associated matrices. The projection matrix is defined as **P**= **V**^-^ - **V**^-^(**X**’**V**^-^**X**)^-^**X**’**V**^-^. The residual maximum likelihood (REML) estimates are obtained using the following equation:

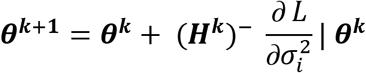

Where, **θ** is the vector of variance components for random effects and covariance components among traits, **H**^-^ is the inverse of the Hessian matrix of second derivatives for the k^th^ cycle, ∂L/σ^2^_i_ is the vector of first derivatives of the likelihood with respect to the variance-covariance components.

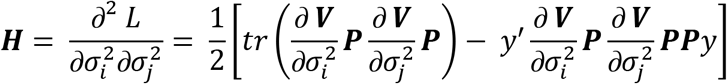

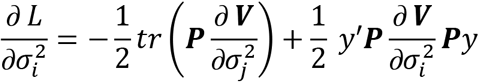

The Eigen decomposition of the relationship matrix proposed by Lee and Van Der Werf (2016) was included in the Newton-Raphson algorithm to improve time efficiency.

### Pin function

Additionally, the pin function to estimate standard errors for linear combinations of variance components (i.e. heritabilities and genetic correlations) was added to the package as well. It uses the delta method to get a first-order approximate standard error for a nonlinear function of a vector of random variables with known or estimated covariance matrix. Suppose *x* is a random vector of length *p* that is at least approximately normally distributed with mean *β* and estimated covariance matrix *C*. Then any function *g*(*β*) of *β*, is estimated by *g*(*x*), which is in large samples normally distributed with mean *g*(*β*) and estimated variance *h*′*Ch*, where *h* is the first derivative of *g*(*β*) with respect to evaluated at *x*. In this setting, *β* is the vector of variance components and *C* is to covariance matrix for the variance components (Fisher’s Information matrix), *g*(*β*) is the function specified by the user, *h* is the first derivative of *g*(*β*) with respect to each variance component involved in the function. The pin function returns both *g*(*x*) and its standard error, the square root of the estimated variance.

### Data sets for performance comparisons

#### BTdata

An ecology dataset for blue tit birds published by Hadfield and colleges (2007), with variables such as tarsus length (tarsus) and colour (back) measured on 828 individuals. The mother of each is also recorded (dam) together with the foster nest (fosternest) in which the chicks were reared. The date on which the first egg in each nest hatched (hatchdate) is recorded together with the sex (sex) of the individuals. This data set was originally used to estimate genetic correlations and heritabilities for behavioural and life-history traits. The model fitted considered tarsus and back as the multivariate response as a function dam and fosternet as random effects with an unstructured covariance structure among traits for such effects. Residuals were also fitted with an unstructured covariance matrix. Sex was fitted as fixed effect.

#### Gryphon data

This is a simulated dataset that was included in the Journal of animal ecology by Wilson and collegues (2010) to help users understand how to use mixed models with animal datasets and pedigree data. Variables indicating the animal (ANIMAL), the mother of the animal (MOTHER), sex of the animal (SEX), and two quantitative traits; birth weight (BWT) and tarsus length at fledging (TARSUS) are available for testing the bivariate model. The model fitted considered TARSUS and BWT as the multivariate response as a function ANIMAL and MOTHER as random effects with an unstructured covariance structure among traits for such effects. Residuals were also fitted with an unstructured covariance matrix. SEX was fitted as fixed effect.

#### CPdata

An outcross (CP) population or F1 cross (between 2 highly heterozygote individuals) in American cranberry (*Vaccinium macrocarpon*) published by Covarrubias-Pazaran and colleges (2017). This dataset contains one year of phenotypic data for 363 siblings. Three phenotypic traits are measured; color (color), yield (Yield), and average fruit weight (FruitAver) and covariates such as row (Row) and column (Col) field information, and genotype (id). About 3000 SNP markers are available for this 363 sibling published by Covarrubias-Pazaran et al. (2016). The model fitted considered color and Yield as the multivariate response as a function id as random effect with an unstructured covariance structure among traits for such effects. Residuals were also fitted with an unstructured covariance matrix.

## Results and Discussion of case examples

### Specification of the model

In order to provide users maximum flexibility to fit heterogeneous-variance and multivariate models, we decided to use a similar formula specification to *ASReml-R* with the *mmer2* function, which works in a formula-based fashion, whereas we kept the *mmer* function as a matrix-based specification where the user provides the incidence and covariance matrices. The different covariance structures available for the random effects and for the trait covariances are show in table 1.

**Table 1.** Different variance-covariance structures available for random effects [var(u)] and variance-covariance structures for traits at different random effects.

| *Random effect variance - $\text{var}(u_i) = \mathbf{G}_i \otimes \mathbf{K}$* |  |  |
| --- | --- | --- |
| $\text{us}(X^*)$ | $\text{diag}(X^*)$ | $\text{at}(X^*)$ |
| $\text{var}(u_i) = \begin{bmatrix} \sigma_{u_1}^2 & \cdots & \sigma_{u_{1,i}} \\ \vdots & \ddots & \vdots \\ \sigma_{u_{1,i}} & \cdots & \sigma_{u_i}^2 \end{bmatrix} \otimes \mathbf{K}$ | $\text{var}(u_i) = \begin{bmatrix} \sigma_{u_1}^2 & \cdots & 0 \\ \vdots & \ddots & \vdots \\ 0 & \cdots & \sigma_{u_i}^2 \end{bmatrix} \otimes \mathbf{K}$ | $\text{var}(u_i) = \begin{bmatrix} \sigma_{u_1}^2 & \cdots & 0 \\ \vdots & \ddots & \vdots \\ 0 & \cdots & \sigma_{u_i}^2 \end{bmatrix} \otimes \mathbf{K}$ |
| Example:<br>`random=~us(ENV):GENO` | Example:<br>`random=~diag(ENV):GENO` | Example:<br>`random=~at(ENV,c(...)):BLOCK`<br>`rcov=~at(ENV,c(...)):units` |

| *Trait variance ($\Sigma$) for random effect and residual specifications* |  |
| --- | --- |
| $\text{us}(\text{trait})$ | $\text{diag}(\text{trait})$ |
| $\begin{bmatrix} \sigma_{t_1}^2 & \cdots & \sigma_{u_{1,i}} \\ \vdots & \ddots & \vdots \\ \sigma_{u_{1,i}} & \cdots & \sigma_{t_i}^2 \end{bmatrix} \otimes \text{var}(u_i)$ | $\begin{bmatrix} \sigma_{t_1}^2 & \cdots & 0 \\ \vdots & \ddots & \vdots \\ 0 & \cdots & \sigma_{t_i}^2 \end{bmatrix} \otimes \text{var}(u_i)$ |
| Examples:<br>`random=~us(trait):us(ENV):GENO`<br>`random=~us(trait):diag(ENV):GENO`<br>`random=~us(trait):GENO`<br>`random=~us(trait):at(ENV):BLOCK`<br>`rcov=~us(trait):at(ENV):units`<br>`rcov=~us(trait):units` | Example:<br>`random=~diag(trait):us(ENV):GENO`<br>`random=~diag(trait):diag(ENV):GENO`<br>`random=~diag(trait):GENO`<br>`random=~diag(trait):at(ENV):BLOCK`<br>`rcov=~diag(trait):at(ENV):units`<br>`rcov=~diag(trait):units` |

\*X refers to any name for a random effect provide by the user and linked to a column name in the data frame provided.
#K refers to any variance covariance matrix for such random effect.

Notice in Table 1 that the *random* and *rcov* arguments allow the user specify different variance functions that provides the flexibility fit some heterogeneous variance models in the **G**_i_ component, i.e.; *at()*, *diag()*, *us()*. Additionally, covariance structures for the covariance among levels of the random effect (**K**) can be provided with the *g()* function, overlay models among random effects with *overlay()*, and customized random effects with the *grp()* function. Detailed documentation in how to use these functions can be found in the help pages within the software (i.e. typing *?grp*). The major enhancement to the *sommer* package is enabling the multi-response models. Variance-covariance structures among traits for the random effects or residuals can be specified with the functionalities *diag(trait)* and *us(trait)* as indicated in Table1.

In order to validate the multivariate version of *sommer* we compared the results and performance of *sommer* with the two most popular multivariate mixed model solvers available in R; *ASReml-R* and *MCMCglmm*.

### BTdata

We fitted a bivariate model for tarsus length (tarsus) and colour (back) as a linear function of the dam and fosternet considering an unstructured covariance structure among traits for both random effects and sex as a fixed effect. When fitting the model using the three software packages we found all provided the same results for variance-covariance components but with differences in time. For example, *ASReml-R* was the fastest, followed by *MCMCglmm* and *sommer* at the end (Table 1). This is due to the nature of the data that has less effects to estimate than observation (n > p) where MME-based algorithms perform faster than direct-inversion algorithms like the one *sommer* uses. When comparing the BLUPs among software (or posterior for *MCMCglmm*) we found a correlation close to 1 showing the equivalence of the software for fitting these models. Standard errors and predicted error variances followed the same trend. Additionally, we used the pin() function to calculate standard errors for the heritabilities and genetic correlations for the *sommer* models to show the capabilities of the package to provide standard errors for linear combination of variance components.

**Table 2.** Comparison of variance components and their standard errors among commonly used R packages for multivariate models (*ASReml-R* and *MCMCglmm*) with *sommer* using the BTdata.

### Gryphon data

We used the simulated dataset from Wilson and collegues (2010) known as the gryphon data (in reference to the mythical animal) to fit a bivariate model for two quantitative traits; birth weight (BWT) and tarsus length at fledging (TARSUS) explained as a linear function of sex (fixed effect) and animal (random effect) assuming an unstructured covariance structured among traits and a covariance structure for the animal random effect provided by the pedigree known as the additive relationship matrix (A). We found *sommer* to provide the same results than *ASReml-R* and *MCMCglmm*. Variance components and their standard errors, along with BLUPs and BLUEs were the same in all software. The only difference was the time for running the model. For example, ASReml-R was the fastest, followed by *MCMCglmm* and *sommer* at the end (Table 2).

**Table 3.** Comparison of variance components and their standard errors among commonly used R packages for multivariate models (*ASReml-R* and *MCMCglmm*) with *sommer* using the gryphon data.

### CPdata

An outcross (CP) population or F1 cross (between 2 highly heterozygote individuals) in American cranberry (*Vaccinium macrocarpon*) published by Covarrubias-Pazaran and colleges (2017) with phenotype and genotype data was used for fitting a multivariate GBLUP model. This dataset contains one year of phenotypic data for 363 siblings. Three phenotypic traits are measured; color, yield, and fruit average weight. We fitted color and yield as the multivariate response as a function of the genotype random effect with a covariance structure that uses the genomic relationship matrix (GBLUP). We found *sommer* to provide the same results than *ASReml-R* and *MCMCglmm*. Variance components and their standard errors, along with BLUPs and BLUEs were the same in all software. This time, *sommer* performed faster than *ASReml-R* and *MCMCglmm* (Table 4).

**Table 4.** Comparison of variance components and their standard errors among commonly used R packages for multivariate models (*ASReml-R* and *MCMCglmm*) with *sommer* using the CPdata.

## Conclusions

We have made available an updated version of *sommer*; a powerful multivariate mixed model solver that allows the fit of heterogeneous variance models with different unknown and known covariance structures based on the Newton Raphson optimization method using direct inversion (DI), which provides a better performance in terms of speed than the MME-based optimization methods when the number of effects to estimate is greater than the number of observations (p > n). In addition, we have made available additional functions for breeding and genetic analysis such as the pin function to estimate standard errors of linear combinations of variance components using the delta method and other functionalities such as two-dimensional splines proposed for modeling the spatial variation of field trials. In a comparison with other software such as ASReml-R, breedR and MCMCglmm (in the univariate/multivariate scenario), *sommer* provided the same results although slower for scenarios of the type p < n and better performance in the p > n scenario. With this effort, we hope to provide tools to researchers to allow them to fit more sophisticated models that reflect better the reality and test their hypothesis with more accuracy.

---

Figure 1. Comparison of BLUPs obtained by sommer, ASReml-R and MCMCglmm in three different datasets (BTdata, gryphon data, CPdata).

